# Phage origin of mitochondrion-localized family A DNA polymerases in kinetoplastids and diplonemids

**DOI:** 10.1101/2020.09.26.314351

**Authors:** Ryo Harada, Yuji Inagaki

**Affiliations:** Graduate School of Life and Environmental Sciences, University of Tsukuba, Japan; Center for Computational Sciences, University of Tsukuba, Japan

**Keywords:** DNA replication, DNA repair, autographivirus, Euglenozoa, lateral gene transfer, mitochondria

## Abstract

Mitochondria retain their own genomes as other bacterial endosymbiont-derived organelles. Nevertheless, no protein for DNA replication and repair is encoded in any mitochondrial genomes (mtDNAs) assessed to date, suggesting the nucleus primarily governs the maintenance of mtDNA. As the proteins of diverse evolutionary origins occupy a large proportion of the current mitochondrial proteomes, we anticipate finding the same evolutionary trend in the nucleus-encoded machinery for mtDNA maintenance. Indeed, none of the DNA polymerases (DNAPs) in the mitochondrial endosymbiont, a putative α-proteobacterium, seemingly had been inherited by their descendants (mitochondria), as none of the known types of mitochondrion-localized DNAP showed a specific affinity to the α-proteobacterial DNAPs. Nevertheless, we currently have no concrete idea of how and when the known types of mitochondrion-localized DNAPs emerged. We here explored the origins of mitochondrion-localized DNAPs after the improvement of the samplings of DNAPs from bacteria and phages/viruses. Past studies revealed that a set of mitochondrion-localized DNAPs in kinetoplastids and diplonemids, namely PolIB, PolIC, PolID, PolI-Perk1/2, and PolI-dipl (henceforth designated collectively as “PolIBCD+”) have emerged from a single DNAP. In this study, we recovered an intimate connection between PolIBCD+ and the DNAPs found in a particular group of phages. Thus, the common ancestor of kinetoplastids and diplonemids most likely converted a laterally acquired phage DNAP into a mitochondrion-localized DNAP that was ancestral to PolIBCD+. The phage origin of PolIBCD+ hints at a potentially large contribution of proteins acquired via non-vertical processes to the machinery for mtDNA maintenance in kinetoplastids and diplonemids.

## Introduction

Mitochondria in the extant eukaryotes are the descendants of an endosymbiotic α-proteobacterium in the last eukaryotic common ancestor (Roger et al. 2017). The mitochondrial (mt) proteins, which are localized in mitochondria, are almost entirely nucleus-encoded and evolutionarily multifarious (Gabaldón and Huynen 2007; Wang and Wu 2014; Gray 2015). Only 10-20% of mt proteins were predicted to be of the α-proteobacterial origin, suggesting that the original proteome of the mitochondrial endosymbiont has been remodeled largely (Gray 2015). There are three possible evolutionary paths that coopt non-α-proteobacterial proteins into the molecular machinery in mitochondria. Non-α-proteobacterial mt proteins could emerge (i) de novo, (ii) by recycling of the pre-existing eukaryotic proteins, or (iii) via lateral gene transfer. Mitochondria, in principle, retain their own genomes that have been descended from the mitochondrial endosymbiont, albeit the entire set of proteins required for mtDNA maintenance (replication and repair) is nucleus-encoded. Thus, as a part of the mitochondrial proteome, the machinery for mtDNA maintenance may be dominated by non-α-proteobacterial proteins. Indeed, none of the known DNA polymerases (DNAPs) localized in mitochondria is most unlikely the direct descendant of the DNAPs in the α-proteobacterial endosymbiont that gave rise to the ancestral mitochondrion (see below).

Phylogenetically diverse eukaryotes possess family A (famA) DNAPs that are evolutionarily related to DNA polymerase I (PolI) in bacteria (Jung et al. 1987; Moriyama et al. 2011). Some of famA DNAPs in eukaryotes are known to be localized in mitochondria (Krasich and Copeland 2017). So far, four distinct types of mitochondrion-localized famA DNAP have been identified. First, “plant and protist organellar DNA polymerase (POP)” appeared to be broadly distributed among eukaryotes (Moriyama et al. 2011; Hirakawa and Watanabe 2019). Second, animals and fungi are known to use DNA polymerase gamma (Polγ) for mtDNA maintenance (Graziewicz et al. 2006). The third type of mitochondrion-localized famA DNAP is “PolIA” shared among members of the classes Kinetoplastea, Diplonemea, and Euglenida, which comprise the phylum Euglenozoa (Klingbeil et al. 2002; Harada et al. 2020). Members of Kinetoplastea and Diplonemea possess the fourth type of mitochondrion-localized famA DNAP. “PolIB,” “PolIC,” and “PolID” were reported originally from a model kinetoplastid *Trypanosoma brucei*, and later identified in broad members of Kinetoplastea (Klingbeil et al. 2002; Harada et al. 2020). The three DNAPs were shown to be closely related to one another in phylogenetic analyses. A recent study further identified multiple DNAPs, which are closely related to but distinct from PolIB, C, or D, in an early-branching kinetoplastid *Perkinsela* sp. and diverse diplonemids (PolI-Perk1/2 and PolI-dipl; Harada et al. 2020). PolIB, C, D, and their related DNAPs were derived from a single molecule, and thus can be regarded collectively as the fourth type of mitochondrion-localized famA DNAP (henceforth termed as “PolIBCD+” in this study). Pioneering studies considered none of the known mitochondrion-localized famA DNAPs as the direct descendant of PolI in the mitochondrial endosymbiont, but failed to clarify how and when POP, Polγ, PolIA, and PolIBCD+ were established in eukaryotic evolution (Moriyama et al. 2011; Hirakawa and Watanabe 2019; Harada et al. 2020).

In this study, we explored the origins of mitochondrion-localized famA DNAPs by analyzing an improved dataset wherein sequence sampling from bacteria and phages was improved drastically. We recovered the intimate affinity between PolIBCD+ and the famA DNAPs of a particular group of phages in phylogenetic analyses. Furthermore, these DNAPs appeared to share a unique insertion of consecutive 8 amino acid (aa) residues. Altogether, we conclude that the extent DNAPs belonging to PolIBCD+ were derived from a single phage famA DNAP acquired by the common ancestor of Kinetoplastea and Diplonemea. We also propose that PolIA in Euglenozoa emerged from a type of cytosolic famA DNAP (Polθ). The origins of PolIA and PolIBCD+ maybe a tip of the remodeling of the machinery of mtDNA maintenance undergone in Kinetoplastea and Diplonemea.

## Results

Prior to this study, the origin of none of the four types of mitochondrion-localized famA DNAPs (i.e. Polγ, POP, PolIA, and PolIBCD+) has been elucidated in detail. This study successfully clarified the origin of PolIBCD+ by analyzing phylogenetic alignments that are much richer in bacterial and phage famA DNAPs than those analyzed in the past studies. The sampling of the bacterial homologs was insufficient to reflect the diversity of bacteria in the previously published phylogenies of famA DNAPs (Moriyama et al. 2011; Hirakawa and Watanabe 2019; Harada et al. 2020). Furthermore, only a few famA DNAPs of phages have been included in the phylogenetic analyses. In this study we prepared the “global famA DNAP” alignment by incorporating diverse bacterial and phage sequences (446 in total) deposited in public databases and 27 sequences that represent the four mitochondrion-localized types of DNAPs (Polγ, POP, PolIA, and PolIBCD+), a single cytosolic DNAP (Polθ), and a single plastid-localized DNAP found exclusively in apicomplexans and chrompodellids (PREX).

The global famA DNAP phylogeny reconstructed four clades, all comprising the eukaryotic homologs exclusively: (i) POP, (ii) PolIA plus Polθ, (iii) PREX, and (iv) Polγ (Shaded in blue in Fig. 1A; see the supplementary materials for the tree with sequence names). The maximum likelihood bootstrap values (MLBPs) for the four clades varied between from 69 to 100%. The POP, PolIA plus Polθ, or Polγ sequences showed no clear affinity to any bacterial or phage famA DNAPs, leaving their origins uncertain. The PREX sequences grouped with bifunctional 3’-5’ exonuclease/DNA polymerases in phylogenetically limited bacteria as previously reported (Janouškovec et al. 2015; Hirakawa and Watanabe 2019; Harada et al. 2020). Curiously, the PolIBCD+ sequences were paraphyletic but nested within a robustly supported clade mainly comprising famA DNAP homologs of phages belonging to families Autographiviridae and Podoviridae (Fig. 1B; this figure corresponds to the portion shaded in gray in Fig. 1A). The famA DNAP homologs of autographiviruses and three bacteria formed a subclade with an MLBP of 72%. The coding regions of two out of the three bacterial famA DNAP homologs in this subclade (marked by stars in Fig. 1B) are flanked by phage-like open reading frames (ORFs) in the corresponding genome assemblies deposited under the GenBank accession Nos LEDQ01000001.1 and NZ_LLYA01000167.1. Phage-like ORFs including that of famA DNAP encompass >40 Kbp consecutively in the two bacterial genomes. Thus, the two “bacterial famA DNAPs” are most likely of lysogenic autographiviruses in bacterial genomes. On the other hand, no phage-like ORF was found around that of famA DNAP in the genome of *Bordetella* genomosp. 9 strain AU14267 (NZ_CP021109.1), suggesting that this bacterium acquired a famA DNAP gene from an autographivirus horizontally. The four PolIBCD+ sequences were positioned at the base of the Autographiviridae clade described above and the grouping of PolIBCD+ sequences and autographivirus famA DNAPs as a whole received an MLBP of 99% (Fig. 1B). The global famA DNAP phylogeny strongly suggests an intimate evolutionary affinity between PolIBCD+ and autographivirus famA DNAPs.

**Fig. 1.**
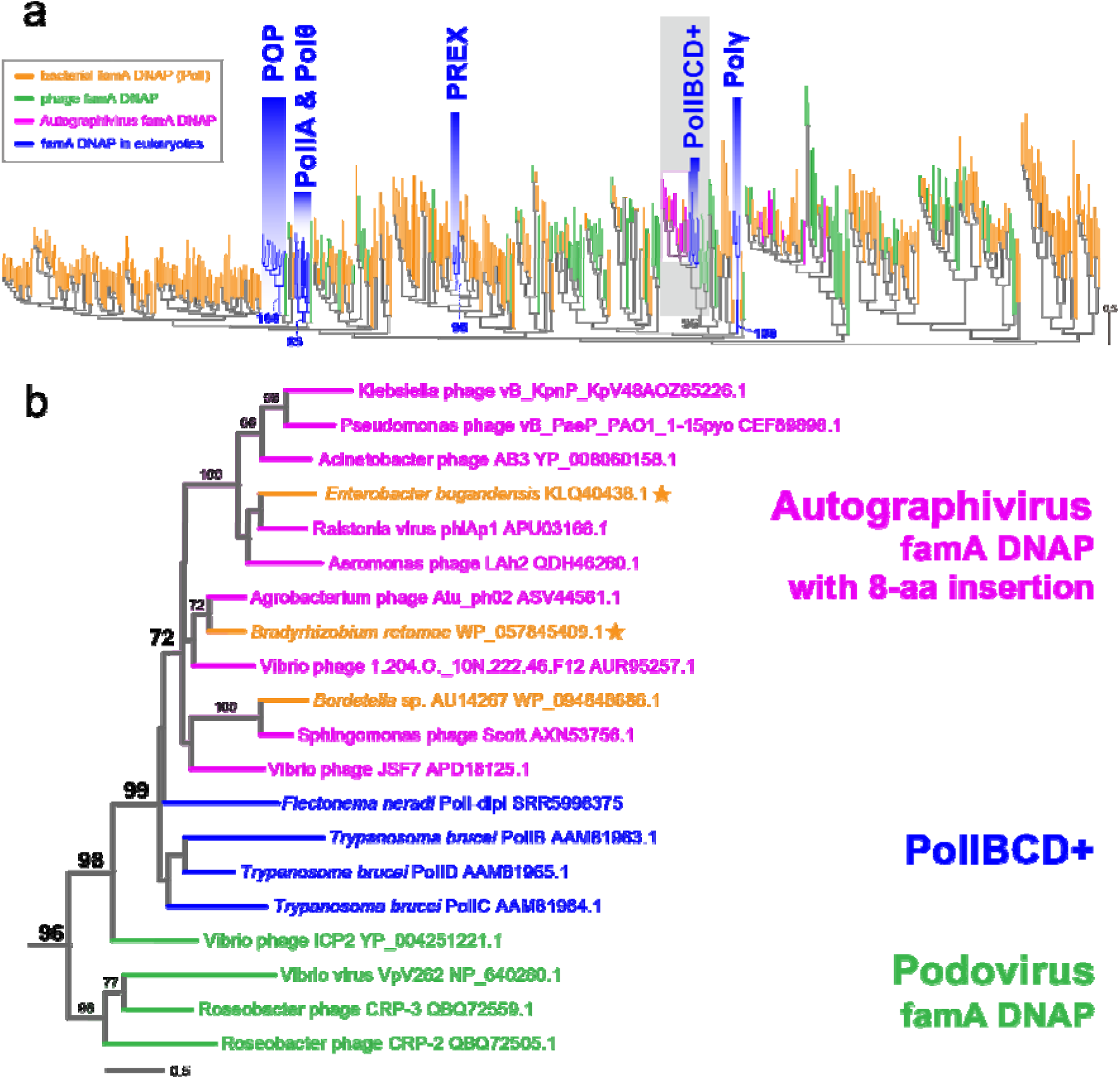
Maximum likelihood (ML) phylogenetic tree inferred from an alignment of the famA DNAP sequences of bacteria, phages/viruses, and eukaryotes. (A) Overview of the entire ML tree. All of the sequence names are omitted. The bacterial and eukaryotic sequences are shown in orange and blue, respectively. The sequences of autographiviruses are shown in magenta. A subset of autographiviruses possess famA DNAPs in the pink-shaded clade bears the characteristic insertion of 8 amino acid residues (AGV^+ins^ famA DNAPs; see the main text for the details). Other phage/viral sequences are shown in green. Only ML bootstrap values of interest are shown. The subtree containing PolIBCD+ and AGV^+ins^ famA DNAP sequences (shaded in gray) is enlarged and presented as (B). ML bootstrap values greater than 70% are shown. AGV^+ins^ famA DNAP sequences marked by stars are of the putative lysogenic phages in bacterial genomes.

Members of Autographiviridae commonly display head-to-tail capsid structures and possess double-stranded linear DNA genomes of approximately 41 Kbp in length. This viral family comprises 9 subfamilies and 132 genera (Lavigne et al. 2008; Adriaenssens et al. 2020). We searched for autographivirus famA DNAPs in the GenBank nr database and detected 175 homologs of 99 members belonging to 57 genera and 76 unclassified members. Each of the 175 members of Autographiviridae seemingly possesses a single famA DNAP. Intriguingly, the autographivirus famA DNAPs were split into two types based on the presence/absence of “8-aa insertion” in the polymerase domain (Fig. S1 and Table S1). In this study, we designate autographivirus famA DNAPs with 8-aa insertion as “AGV^+ins^ famA DNAPs”. Each AGV^+ins^ famA DNAPs was predicted to possess only polymerase domain by InterProScan5 with the Pfam database (Jones et al. 2014; El-Gebali et al. 2019) (Table. S2). AGV^+ins^ famA DNAPs were found in 40 members belonging to 23 genera, and 51 unclassified members (Fig. S1). Although only a subset of the 175 autographivirus famA DNAPs was included, the global famA DNAP phylogeny (Fig. 1A) demonstrated the distant relationship between AGV^+ins^ famA DNAPs and other autographivirus famA DNAPs lacking 8-aa insertions.

To reexamine the phylogenetic affinity between PolIBCD+ and AGV^+ins^ famA DNAPs, we selected non-redundant sequences from the 91 AGV^+ins^ famA DNAPs and aligned with 24 PolIBCD+ sequences and four famA DNAPs of phages belonging to a family Podoviridae as the outgroup. The second famA DNAP alignment was subjected to both ML and Bayesian methods. In the second phylogenetic analyses, AGV^+ins^ famA DNAPs and PolIBCD+ sequences formed a clade supported by an MLBP of 100% and a Bayesian posterior probability (BPP) of 1.0 (Fig. 2). PolIBCD+ sequences appeared to possess 8 amino acids that are most likely homologous to 8-aa insertion in AGV^+ins^ famA DNAPs (Fig. 2), strengthening the phylogenetic affinity between PolIBCD+ and AGV^+ins^ famA DNAPs. Besides PolIBCD+ and AGV^+ins^ famA DNAPs, 8-aa insertion was found solely in the famA DNAP homolog of Vibrio phage ICP2 placed at the basal position of the clade of PolIBCD+ and AGV^+ins^ famA DNAPs (Fig. 2). In the analyses of the second alignment, AGV^+ins^ famA DNAPs grouped together with an MLBP of 92% and a BPP of 0.99, excluding PolIBCD+ sequences that formed a clade with an MLBP of 72% and a BPP of 0.66 (Fig. 2). The weak statistical support for the monophyly of PolIBCD+ sequences is not incongruent with their paraphyletic relationship reconstructed in the global famA DNAP analysis (Fig. 1B).

**Fig. 2.**
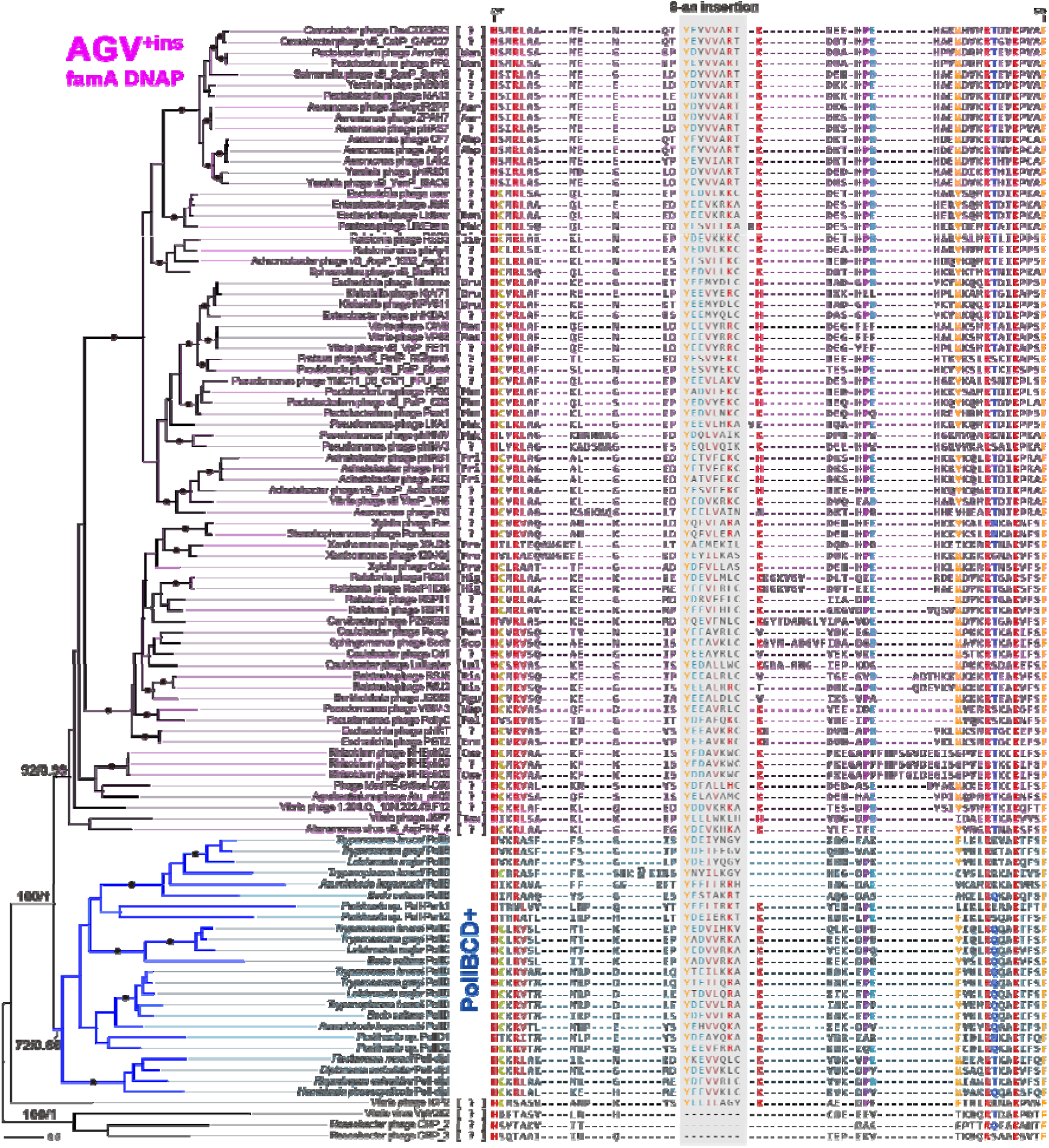
Phylogenetic relationship among 74 AGV^+ins^ famA DNAP and 24 PolIBCD+ sequences that share a unique insertion of 8 amino acid residues (8-aa insertion). The tree topology and branch lengths inferred by the maximum likelihood (ML) method are shown on the left. ML bootstrap values (MLBPs) and Bayesian posterior probabilities (BPPs) for only the nodes critical to infer the origin of PolIBCD+ are shown. As ML and Bayesian analyses reconstructed the essentially same tree topology, only BPPs for the selected nodes are presented. The nodes supported by an MLBP of 100% and a BPP of 1.0 are marked by dots. The genus names of the autographiviruses (and podoviruses), from which famA DNAPs were sampled, are given in brackets. Abbreviations are follows: Aer, Aerosvirus; Ahp, Ahphunavirus; Bon, Bonnellvirus; Cue, Cuernavacavirus; Dru, Drulisvirus; Erm, Ermolevavirus; Fri, Friunavirus; Hig, Higashivirus; Jia, Jiaoyazivirus; Kal, Kalppathivirus; Lul, Lullwatervirus; Mac, Maculvirus; Mgu, Mguuvirus; Nap, Napahaivirus; Per, Percyvirus; Phk, Phikmvvirus; Phm, Phimunavirus; Pol, Pollyceevirus; Pra, Pradovirus; Ris, Risjevirus; Sco, Scottvirus; Taw, Tawavirus; Wan, Wanjuvirus; ?, unclassified. The amino acid sequences of 8-aa insertions and their flanking regions are shown on the right. 8-aa insertions are shaded in grey. The residues are colored according to their degrees of conservation. The amino acid residue numbers shown on the left and right edges of the alignment are based on the famA DNAPs of Cronobacter phage DevCD23823 (YP_009223394.1).

## Discussion

The phylogenetic analyses of the global alignment of famA DNAPs (Figs. 1A and 1B) and the second alignment rich in AGV^+ins^ famA DNAP homologs (Fig. 2) consistently recovered the specific affinity between PolIBCD+ and AGV^+ins^ famA DNAPs. These results strongly suggest that PolIBCD+ in the extant kinetoplastids and diplonemids can be traced back to a single autographivirus famA DNAP, particularly the one with 8-aa insertion. In other words, PolIBCD+ is a typical example of non-α-proteobacterial mt proteins established via lateral gene transfer. Unfortunately, even the analyses of the second alignment, wherein the known diversity of AGV^+ins^ famA DNAPs was covered, failed to pinpoint the exact origin of PolIBCD+ (Fig. 2). We might be able to find an AGV^+ins^ famA DNAP homolog that branches PolIBCD+ sequences directly in a future phylogenetic study covering the true diversity of phage famA DNAPs. In particular, we regard that autographivirus famA DNAP genes in bacterial genomes are significant. To our knowledge, no autographivirus has been reported to infect eukaryotes. Thus, the common ancestor of kinetoplastids and diplonemids may have acquired the famA DNAP gene from a lysogenic autographivirus in a bacterial genome. If so, the bacterial genomes harboring AGV^+ins^ famA DNAP genes are critical to investigate the origin of PolIBCD+ at a finer level than that in the current study.

Members of classes Kinetoplastea and Diplonemea, together with Euglenida, share another type of mitochondrion-localized famA DNAP, namely PolIA (Harada et al. 2020). It is reasonable to postulate that the common ancestor of the three classes—most likely the ancestral euglenozoan—had established the ancestral PolIA. Although the origin of PolIA has not been addressed explicitly, past studies recovered the phylogenetic link between PolIA and Polθ, a type of famA DNAP operated in the cytosol of eukaryotic cells. The original study reporting PolIA, B, C, and D in *Trypanosoma brucei* has hinted at the phylogenetic affinity between PolIA and Polθ (Klingbeil et al. 2002). A recent phylogeny including famA DNAPs sampled from eukaryotes and limited bacteria (Note that no phage homolog was included) reconstructed a clade of PolIA and Polθ sequences with high statistical support (Harada et al. 2020). The PolIA-Polθ affinity persisted even after the sampling of famA DNAPs from bacteria and phages was improved drastically in this study (Fig. 1A). We here propose that the ancestral PolIA was likely derived from a Polθ homolog followed by the change in subcellular localization from the cytosol to the mitochondrion. Noteworthy, the evolutionary processes yielded PolIA and PolIBCD+, both of which are mt proteins of non-α-proteobacterial origin, are different substantially from each other. The former emerged through the recycling of a pre-existing eukaryotic protein while the latter is of phage origin (See above). The Polθ origin of PolIA is the best estimate from both past and current phylogenetic analyses of famA DNAPs but alternative possibilities still need to be explored in future studies.

The repertories of mitochondrion-localized DNAPs in euglenozoans appeared to be more complex than those in the majority of other eukaryotes in which a single type of mitochondrion-localized DNAP (i.e. POP or Polγ) seemingly operates. The complexity in the repertory of DNAPs in euglenozoan mitochondria seems to coincide with that in the structure of their mtDNAs (Lukeš et al. 2002; Roy et al. 2007; Spencer and Gray 2011; Dobáková et al. 2015; Yabuki et al. 2016; Burger and Valach 2018). Nevertheless, it is unlikely that the non-α-proteobacterial background is restricted to PolIA and PolIBCD+ among the proteins involved in mtDNA maintenance. Rather, the machinery for mtDNA maintenance in the common ancestor of kinetoplastids and diplonemids (and its descendants) are heavily remodeled by both incorporating exogenous proteins via lateral gene transfer and recycling the pre-existed nucleus-encoded proteins. The above conjecture can be examined only after we identify the major proteins involved in DNA maintenance in kinetoplastid/diplonemid mitochondria and their evolutionary origins.

## Materials & Methods

### Global phylogeny of famA DNAPs

We searched for the amino acid (aa) sequences of bacterial and phage famA DNAPs in the NCBI nr database as of March 6, 2020, by BLASTP using the polymerase domain of *Escherichia coli* PolI (KHH06131.1; the portion corresponding to the 491^st^–928^th^ aa residues) as a query (Camacho et al. 2009; Sayers et al. 2020). We retrieved the sequences matched to the query with *E* values equal to or less than 1 × 10^−4^ and covered more than 200 aa in the polymerase domain. Note that the sequences derived from metagenome analyses were excluded from this study. The redundancy within famA DNAP sequences was removed by cluster analysis using CD-HIT v4.7 with a threshold of 40% (Li and Godzik 2006; Fu et al. 2012). We finally selected 119 and 327 aa sequences of phage and bacterial famA DNAPs, respectively, for the downstream analyses (see below).

The bacterial and phage famA DNAP aa sequences (446 in total) were aligned with those in eukaryotes (27 in total), namely (i) mitochondrion-localized famA DNAPs in Kinetoplastea and Diplonemea (PolIA, B, C, D, and PolI-dipl), (ii) mitochondrion-localized famA DNAPs in animals and fungi (Polγ), (iii) Polθ localized in the cytosol, (iv) mitochondrion and/or plastid-localized famA DNAPs in diverse eukaryotes (POP), and (v) plastid-localized famA DNAPs in apicomplexan parasites and their relatives (PREX). The aa sequences were aligned by MAFFT v7.455 with the L-INS-i model (Katoh and Standley 2013). Ambiguously aligned positions were discarded manually, and gap-containing positions were trimmed by using trimAl v1.4 with the -gt 0.95 option (Capella-Gutiérrez et al. 2009). The final “global famA DNAP” alignment comprised 473 sequences with 316 unambiguously aligned aa positions. The final global famA alignment is provided as a part of the supplementary materials. We subjected this alignment to the ML phylogenetic analysis by IQ-TREE v1.6.12 using the LG + Γ + F + C60 + PMSF model (Nguyen et al. 2015; Wang et al. 2018). The guide tree was obtained using the LG + Γ + F model that was selected by ModelFinder (Kalyaanamoorthy et al. 2017). The statistical support for each bipartition in the ML tree was calculated by 100-replicate non-parametric bootstrap analysis.

### Phylogenetic analyses of an alignment rich in autographivirus famA DNAPs

We retrieved 175 famA DNAP aa sequences of autographiviruses from the NCBI nr database. The details of the survey were the same as described above. The 175 famA DNAPs were sampled from 99 members belonging to 57 genera and 76 unclassified members in the family Autographiviridae. The autographivirus famA DNAPs were found to comprise two types based on the presence/absence of an insertion of 8 aa residues (8-aa insertion; see above). The famA DNAPs with 8-aa insertion (AGV^+ins^ famA DNAPs) appeared to be closely related to PolIBCD+, mitochondrion-localized famA DNAPs in kinetoplastids (PolIB, C, D, PolI-Perk1/2) and that in diplonemids (PolI-dipl). The redundancy among the AGV^+ins^ famA DNAPs was reduced by cluster analysis using CD-HIT v4.7 with a threshold of 90%. Finally, we aligned the aa sequences of 74 AGV^+ins^ famA DNAPs, 24 PolIBCD+, and famA DNAPs of four members of Podoviridae by MAFFT v7.455 with the L-INS-i model. Ambiguously aligned positions were discarded manually, and gap-containing positions were trimmed by using trimAl v1.4 with the -gt 0.9 option. The final version of the second alignment is provided as a part of the supplementary materials. The final alignment containing 102 sequences with 581 unambiguously aligned aa positions was subjected to both ML and Bayesian phylogenetic analyses. The ML and ML bootstrap analyses were performed as described above. For Bayesian analysis using Phylobayes v4.1, we run four Markov Chain Monte Carlo chains for 100,000 cycles with burn-in of 25,000 (maxdiff = 0.09472) and calculated the consensus tree with branch lengths and BPPs from the remaining trees (Lartillot et al. 2009). The amino acid substitution model was set to CAT + GTR in Phylobayes analysis described above.

## Data availability

The alignment datasets for phylogenetic analysis are available in supplementary materials at https://drive.google.com/drive/folders/1vpwh0MzYul_wjKmyutZIZR1MSmMZn5ca?usp=sharing.

## Acknowledgments

This work was supported by the grants from the Japanese Society for Promotion of Sciences awarded to Y. I. (numbers 18KK0203 and 19H03280).

